# Scalable variational inference for super resolution microscopy

**DOI:** 10.1101/081703

**Authors:** Ruoxi Sun, Evan Archer, Liam Paninski

## Abstract

Super-resolution microscopy methods (e.g. STORM or PALM imaging) have become essential tools in biology, opening up a variety of new questions that were previously inaccessible with standard light microscopy methods. In this paper we develop new Bayesian image processing methods that extend the reach of super-resolution microscopy even further. Our method couples variational inference techniques with a data summarization based on Laplace approximation to ensure computational scalability. Our formulation makes it straightforward to incorporate prior information about the underlying sample to further improve accuracy. The proposed method obtains dramatic resolution improvements over previous methods while retaining computational tractability.

## Introduction

Super-resolution microscopy techniques, such as STORM [1] or PALM [2] imaging, have quickly become essential tools in biology. These methods overcome the light diffraction barrier of traditional microscopy, thus enabling researchers to ask questions previously considered inaccessible (as a measure of impact, developers of these methods were awarded the Nobel prize in Chemistry in 2014). Given a sample treated with a fluorescent dye, the basic strategy is to stochastically activate fluorophores at a low rate, guaranteeing that only a sparse subset are activated at a given time. By repeatedly imaging the sample we obtain a movie wherein each frame reflects a random, sparse set of fluorophore activations. Then we exploit the sparsity of activations within each frame to localize the positions of the activated fluorophores; aggregating a long sequence of such point localizations then yields a super-resolved image (Fig. 1).

**Figure 1:**
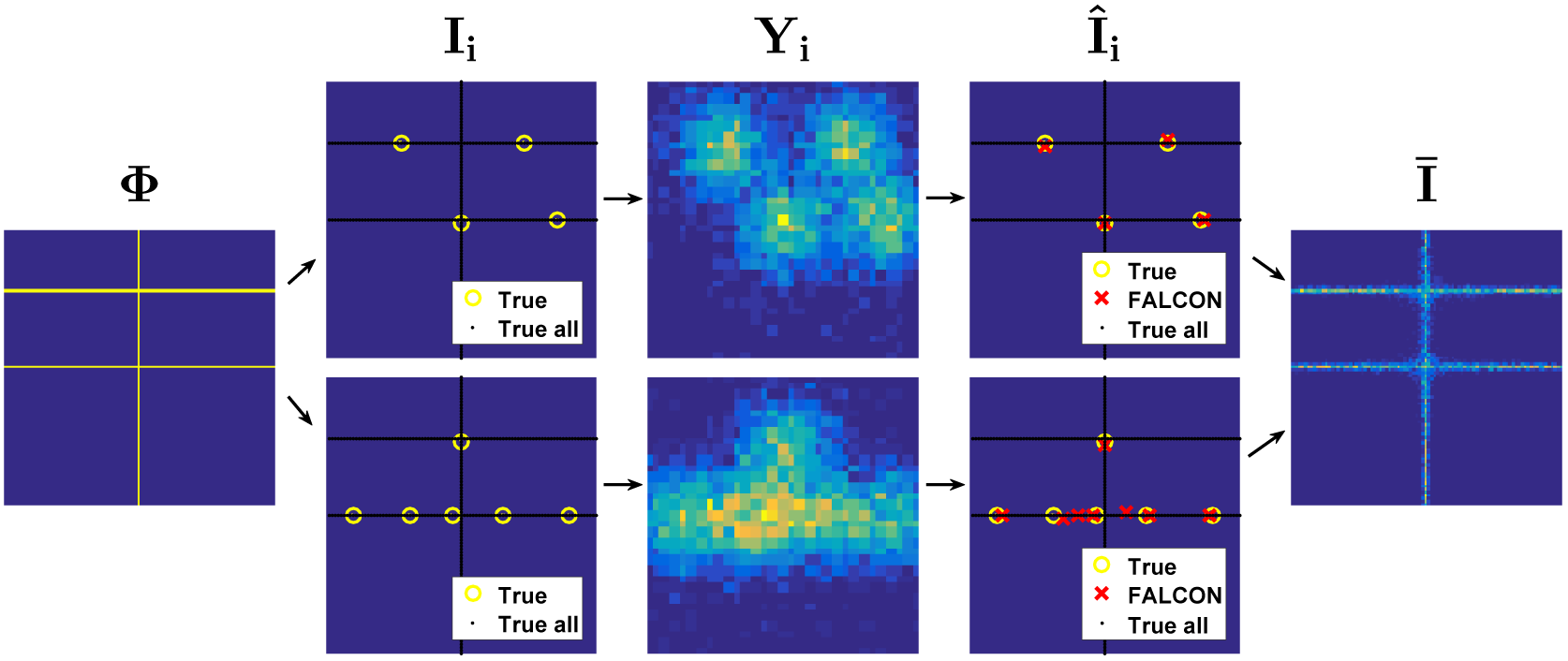
Overview of standard super-resolution microscopy. **Column 1**: The true fluorophore density matrix Φ. **Column 2**: *I*_*i*_ indicates the sparse subset of fluorophores (yellow circles) activated on frame *i* from Φ, which is also plotted as background (black dots); two independent sample frames shown here (top and bottom). **Column 3**: *Y*_*i*_ are the observed camera images on these two frames, formed by blurring and downsampling the corresponding *I*_*i*_ and adding Poisson noise. **Column 4**: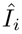 indicate the estimated locations of the active fluorophores on each frame, with the true *I*_*i*_ shown for comparison. The FALCON method [4] was used to compute the estimates here; note that estimator performance decreases in regions where the “bumps” in *Y*_*i*_ overlap significantly. **Column 5**: The standard approach to estimate *p*Φ is to simply average over multiple inferred frames, 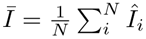.

Many methods have been proposed to expand upon this basic idea, focusing upon improving localization performance within each individual frame [3]. For sparse recovery of a single frame, several modern techniques take a compressed sensing approach that exploits the true sparsity of the underlying fluorophore activations; these techniques result in a formulation as a sparse deconvolution problem [4, 5], providing scalable, fairly accurate reconstructions.

The critical message of this paper is that such standard approaches are sub-optimal because each frame is reconstructed independently, thereby discarding information that should be shared across frames. Intuitively, given *N* – 1 reconstructed frames, we should have a good deal of prior information about the locations of fluorophores on the *N*-th frame, and ignoring this information will in general lead to highly suboptimal estimates. (This basic point has been made previously, e.g. by [6, 7]; we will discuss this work further below.)

Here, we propose a scalable Bayesian approach that properly pools information across frames and can also incorporate prior information about the image, leading to dramatic resolution improvements over previous methods while retaining computational tractability.

## Model

At each frame *i* we observe an *L* × *L* fluorescence image 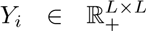, and collect the sequences of *N* observed frames into the movie *Y = {Y_i_*: *i* ∈ {1,…, *N*}}. We model each observed frame *Y_i_* as a noisy, blurred, low-resolution image,

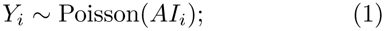

here *A* is a matrix implementing convolution with a known point-spread function (PSF), scaling by the mean photon emission rate per fluorophore, and spatial downsampling; the high-resolution image 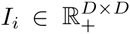 is a sparse matrix, zero except at the locations of fluorophores activated on frame *i.* In this application *L* < *D.* Below we will use the sparse representation (m_i_, *F*_*i*_) for *I*_*i*_: *m*_*i*_ denotes the number of active fluorophores in *I*_*i*_ and 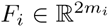 denotes the vector of *xy* positions of these fluorophores. Note that multiple fluorophores can be active at the same location, so the entries of *I*_*i*_ are nonnegative integers; it is straightforward to extend our methods to the case that *I*_*i*_ can take arbitrary nonnegative real values, but we will suppress this case here for notational simplicity.

At each high-resolution pixel position (*x,y*), we model the activation of fluorophores by an inhomogeneous Poisson process with rate *λ*_*xy*_,

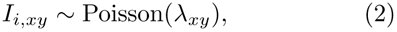

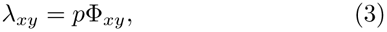

and *p* is a scalar (typically under at least partial experimental control) that sets the fluorophore emission rate. The matrix 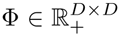 specifies the density of fluorophores at each pixel location, and is the main object we aim to estimate; since λ and Φ are related by a constant (*p*), we will develop the inference methods below in terms of λ, as this leads to slightly simpler algebra.

This model can be extended to 3*D* [8, 9] and/or multispectral imaging [10], but for simplicity here we focus on 2D single-color imaging.

The above definitions lead to the joint probability distribution

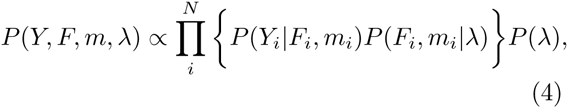

where *m* and *F* collect the *N* scalars *m*_*i*_ and vectors *F*_*i*_, respectively; P(λ) is a prior distribution on λ

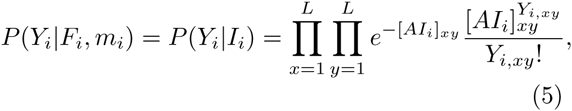

where we have used the equivalence between *I*_*i*_ and (*m*_*i*_ *F*_*i*_), and

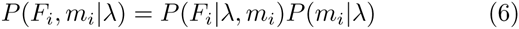

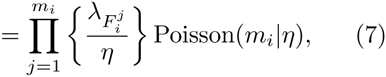

where 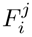 denotes the *xy* position of the *j*-th active fluorophore in frame 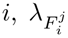 is the value of the 2D function λ at location 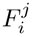, and we have abbreviated the normalizer 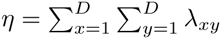

## Inference

Now that the model and likelihoods have been defined, we can proceed to develop our estimator for the underlying fluorophore density image λ. We take a Bayesian approach, which requires that we approximate the posterior distributions of the unknown quantities (*m*_*i*_,*F*_*i*_) given the observed data *Y*_*i*_. (Approximation methods are required here since this is a non-conjugate latent variable model; we cannot analytically integrate out the *I*_*i*_ variables.) A number of such approximation methods are available; for example, [11] recently developed MCMC methods to perform Bayesian inference in a similar model. However, these methods do not scale to the cases of interest here, where the number of frames *N* and pixels (*D*^2^ and *L^2^*) are often quite large.

Therefore we have developed a variational expectation-maximization (vEM) [12] approximate inference approach. As is standard, we need to choose a variational family of distributions *q* (these distributions will be used to approximate the true posterior), then write down the “evidence lower bound” (ELBO; this is a function of *q* and other model parameters), and then develop methods for tractably ascending the ELBO.

The most standard choice of *q* here (a fully factorized distribution over all latent variables, i.e., the activations *I*_*i*_ together with λ) does not lead to a scalable inference method, due to the very high dimensionality of {*I*_*i*_}; in addition, this vanilla variational approximation is poor here because of strong posterior correlations between adjacent pixels in the *I*_*i*_ images. Instead, we exploit the sparse representation (*m*_*i*_, *F*_*i*_) for *I*_*i*_: a more effective approach for approximating *p(F, m|Y*, λ) was to use a simple point estimate for *m* (discussed below) and then, conditionally on *m*, a factorized (“mean-field”) approximation for *p(F|m, Y*, λ). Thus we approximate

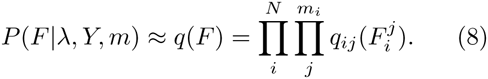

We have factorized across frames *i* and active fluorophores *j* within each frame; here each 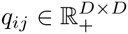 is a probability density on the *D* × *D* grid that summarizes our approximate posterior beliefs about the fluorophore location 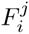. In practice each *q*_*ij*_ will be extremely sparse, with very compact support, as we will discuss further below (Fig 2E).

**Figure 2:**
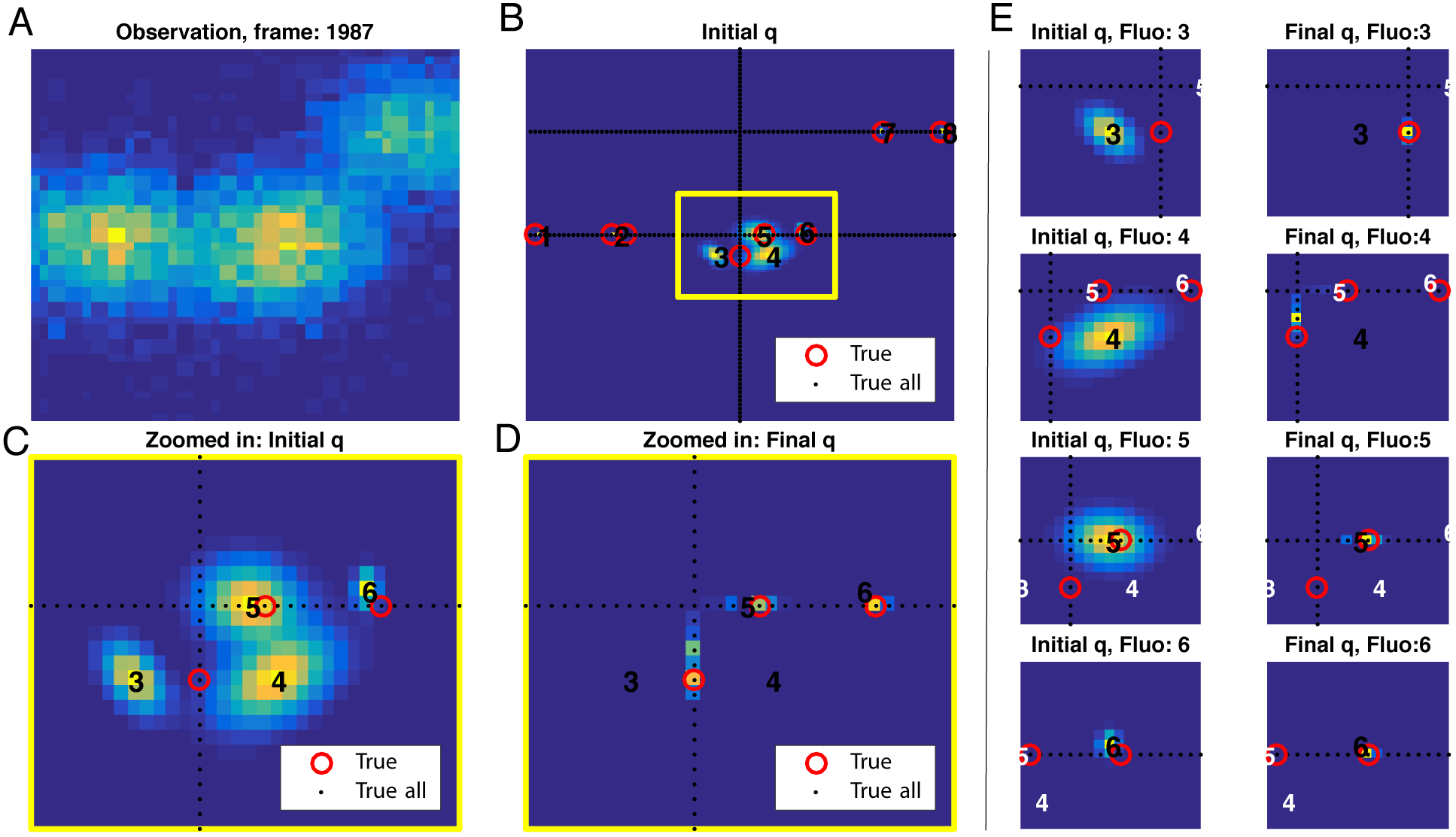
Updating the factors *q*_*ij*_ in the E step. (A) A single simulated observation frame *Y*_*i*_. **(B)** 8 superimposed initial *q*_*ij*_ distributions for frame *i*, computed via Laplace approximations with means 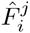. True active fluorophores in *I*_*i*_ are labeled as red circles; true λ indicated by black dots; numbers indicate each 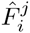 (ordering is arbitrary). Fluorophores 1,2,7, and 8 are relatively spatially isolated, with correspondingly large Fisher information (see supplementary Fig. 8) and so their initial *q* distributions are highly concentrated (and cannot even be seen beneath the red circles). In contrast, the closely overlapping PSF’s of fluorophores 3,4,5, and 6 lead to broad initializations of *q.* **(C)** Zoom of yellow region outlined in **B. (D)** Final *q*_*ij*_’s estimated by the vEM algorithm (same region as in panel **B**). Note that these have converged onto the region of positive λ (despite not having access to the ground truth λ), and the four original estimated fluorophore distributions have essentially converged near the 3 true active fluorophores in this region. **(E)** Further zoom showing details of each *q*_*ij*_ in **D**. The locations of other 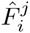 are indicated by white numbers. Left column: initial *q*_*ij*_’s; right column: final *q*_*ij*_’s. Again, the numbers indicate the 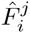 locations, which correspond to the peaks of the initial *q*_*ij*_’s. Note the significant differences between the initial and final *q*_*ij*_’s.

The ELBO is given by:

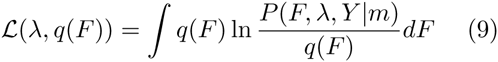

Our goal is to maximize ℒ (*λ, q*(*F*)) with respect to the distributions *q*_*ij*_ and image λ.

We will use a coordinate-ascent approach in which we update one *q*_*ij*_ or λ at a time; as discussed below, after one more approximation each update step can be computed cheaply (and parallelizes easily), and empirically only a few coordinate sweeps are necessary for convergence to a local optimum.

## Laplace Approximation

Computing each *q*_*ij*_ update directly requires the computation of an *L* × *L* sum over the observed data image *Y*_*i*_ and several *D* × *D* sums over the other factors 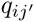, and since we have to compute these updates repeatedly, it is important to reduce the computation time in this inner loop. We have found that we can effectively summarize the data in each frame by using a conditional Laplace approximation to the likelihood. Specifically, we approximate

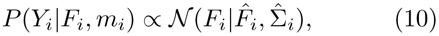

where the left hand side is the Poisson likelihood from eq. 5 and the right hand side denotes a multivariate normal density over 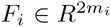, with mean

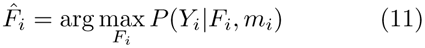

and covariance inverse to the Fisher information *J*_*i*_,

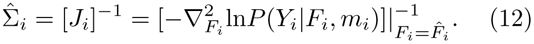

This Gaussian approximation to the Poisson likelihood is well-known to be accurate in the high-information regime where a sufficient number of photons are observed; see [13] for further discussion, and supplementary Fig. 8 for empirical evaluations of this approximation in the context of our simulations. (However, note that this Laplace approximation is not equivalent to assuming a Gaussian noise model with constant variance for *Y*_*i*_; the Poisson noise model used here is significantly more accurate and consistent with the physics of shot noise.)

As we will see in the next subsection, this approximation allows us to replace the expensive sums noted above with evaluations of a much simpler 2*m*_*i*_-dimensional quadratic form. 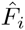 and 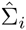 serve as approximate sufficient statistics for *Y*_*i*_, drastically reducing the size of the data that needs to be touched per iteration. In fact, the observed Fisher information matrix *J*_*i*_ is sparse – if fluorophores *j* and *j’* are sufficiently distant (more than a couple PSF widths apart) then 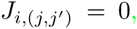, and this can be used to further speed up the computation. In practice, we compute *J*_*i*_ via automatic differentiation [14] and locally optimize eq. 11 numerically using an efficient initializer discussed further below.

The Laplace approximation also provides a convenient initialization for the *q*_*ij*_’s: we simply set each *q*_*ij*_ to be the marginal (Gaussian) density of 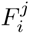 in eq. 10, with *q*_ij_ set to zero for all pixels sufficiently distant from 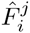.

## Variational EM Algorithm

Now we can put the pieces together and derive our vEM algorithm. The first step is to expand the ELBO eq.9, plugging in our factorized q, the Laplace approximation eq.10, and the likelihood eq.7 to arrive at

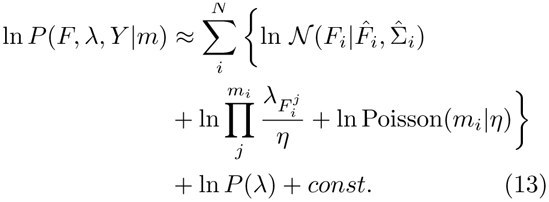

The vEM algorithm alternates between an E step (in which we optimize the ELBO wrt each *q*_*ij*_, with λ and all the other *q*’s held fixed), and an M step (in which we optimize the ELBO wrt λ with all the *q*’s held fixed).

### M step

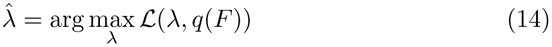

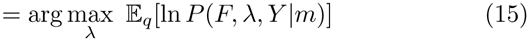

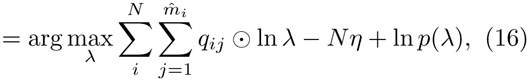

with ⊙ denoting pointwise multiplication. If we use a flat prior for λ, the *p(λ*) term can be dropped, and if we abbreviate 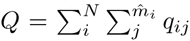, we have the solution 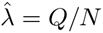. (Recall that 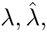 and 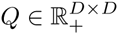 This is a natural generalization of the MLE for a discretized inhomogeneous Poisson process.

### E step

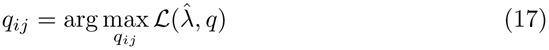

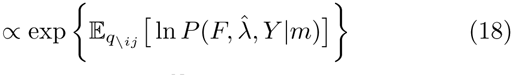

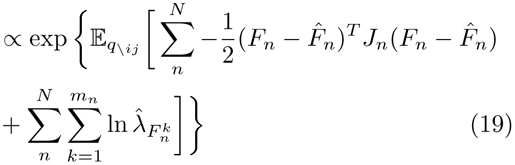

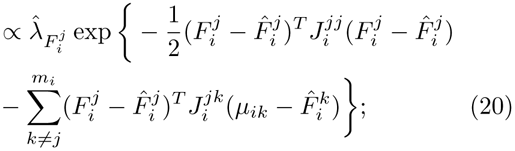

here we have abbreviated 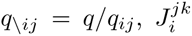 is the 2 × 2 block of *J*_*i*_ corresponding to fluorophores *j* and *k*, and 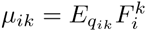. Note that in the end, due to the Laplace approximation, the *q*_*ik*_’s only enter the update above via their means, and that the updated *q*_*ij*_ is simply proportional to a Gaussian factor multiplied by 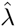.

Finally, note that the effective support of each *q*_*ij*_ tends to shrink compared to the initialization (and this increasing sparsity can be readily exploited computationally); this makes sense, because our initialization (from the Laplace approximation) is based only on the likelihood of a single frame *Y*_*i*_ – when we incorporate the information from other frames (via 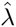) the approximate posterior *q*_*ij*_ tends to become more concentrated. See Fig. 2 for an illustration.

## Extensions and further details

In the developments above we have deferred several questions. How do we estimate *m*_*i*_, the number of active fluorophores in each frame? How do we initialize the optimization problem eq.11 in the Laplace approximation for the likelihood? How do we make use of prior information *P(λ*) in the M-step?

For the first two tasks mentioned above we exploit pre-existing solutions. Specifically, we have found that the FALCON [4] method provides fairly good preliminary estimates of both the number and the location of fluorophores in each frame *i;* the former is used as *m*_*i*_ and the latter are used to initialize the optimization in eq. 11.

One of the major benefits of a Bayesian approach is that we can easily incorporate prior information about parameters of interest – in this case, λ. In principle it is possible to incorporate various sources of prior information about λ, but here we restrict our attention to the simplest case: in many cases the true underlying λ is known to be sparse, and we can exploit this fact to improve our estimates significantly. (Note that this sparsity constraint on λ is in addition to the fact that the images I_i_ are sparse, a fact that we have already exploited repeatedly. Also note that standard super-resolution approaches exploit the sparsity of each *I*_*i*_ – but since they simply average over the estimated 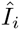 to obtain 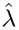, previous approaches have not attempted to further exploit the sparsity of λ.) An effective and computationally trivial approach is to apply the standard L1 “soft threshold” operator [15] to *Q* in the M-step (eq.16):

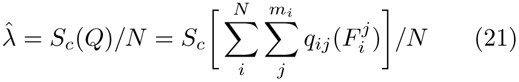

where we define *S*_*c*_[*x*] = max(*x – c*, 0); the threshold *c* can easily be chosen to achieve an a desired level of sparsity (typically set by prior knowledge, though cross-validation could be used here instead).

When active fluorophores are well-isolated in the image (i.e., the “bumps” corresponding to each active fluorophore are sufficiently distinguishable) then FALCON’s estimates are typically accurate, and the corresponding entries of the Fisher information matrix *J*_*i*_ are large. However, when the bumps overlap then the Fisher information can decrease significantly (see supplementary Fig. 8 for an illustration) and the accuracy of m_i_ and the nearby fluorophore estimates 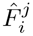 decrease. In this case we can achieve significantly improved accuracy by exploiting information from other frames, via the estimated 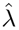.

Thus the full algorithm proceeds as follows. To initialize we run FALCON on each frame and compute the Laplace approximation, then run vEM (restricting attention to the ~ 50% of frames on which fluorophore activation was sparsest, to improve localization accuracy). Then we rerun FALCON incorporating information from the preliminary 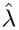 estimate to improve the estimates 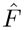 and 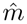. FALCON uses an L1-penalized regression approach to obtain preliminary estimates of *I*_*i*_ from *Y*_*i*_; it is straight forward to include a weighted L1 term where the weight is inversely proportional to 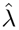 to encourage the FALCON output to localize near regions of high 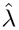 (and to eliminate some spurious location estimates). Then we can use the resulting updated 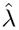-constrained FALCON estimates of the fluorophore locations to re-initialize eq. 11 on the subset of frames where the preliminary FALCON and vEM results disagree (updating these *m*_*i*_ as well), and proceed with further vEM iterations. This procedure can in principle be iterated, though we find in practice that one outer iteration typically suffices. In the inner loop, we found that just 5 vEM iterations were sufficient. See Fig. 3 for an illustration of each algorithm step’s output.

**Figure 3:**
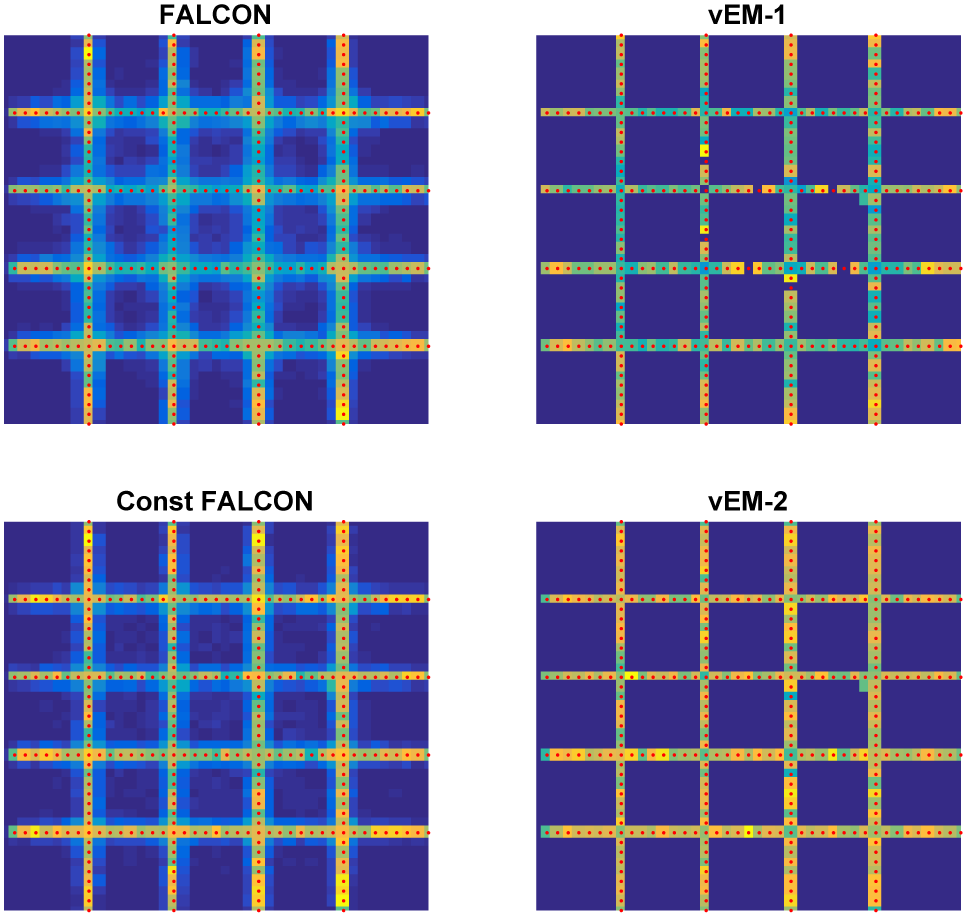
Illustrating the algorithm steps on a simulated example. Upper left: FALCON estimate given 5000 frames of data. Upper right: output after first run of vEM, based on the 2000 sparsest frames. This estimate is used to constrain FALCON to obtain a more accurate preliminary estimate 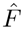 and 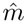, with all 5000 frames (lower left), and our final estimate using all 5000 frames after a second vEM run is shown in the lower right. The grid shape of the true underlying simulated image (red dots) is recovered essentially perfectly in the lower right; estimation noise (averaged over frames) blurs the true grid shape significantly in the left panels.

## Results

Figures 3–6 detail simulated comparisons between FALCON, a state-of-the-art super-resolution algorithm [4], and the vEM algorithm developed here. The simulated image was a simple grid pattern; full simulation details are given in the Appendix. In Fig. 3 it is clear that the variational EM algorithm recovers the true grid support in this simulated example more accurately than does the FALCON algorithm. Fig. 4 quantifies the performance of the new proposed algorithm following each step illustrated in Fig. 3. Specifically, we examine the proportion of fluorophores whose estimated positions were recovered on the correct support of the true underlying grid image (panel B); the frame-by-frame precision and recall (and F measure, defined as the harmonic mean of precision and recall) of individual fluorophore estimates (panel C); and the frame-by-frame absolute error of individual fluorophore estimates (panel D). In each panel, we see that vEM leads to significant improvements over FALCON; applying soft thresholding in the M-step also leads to significant improvements.

**Figure 4:**
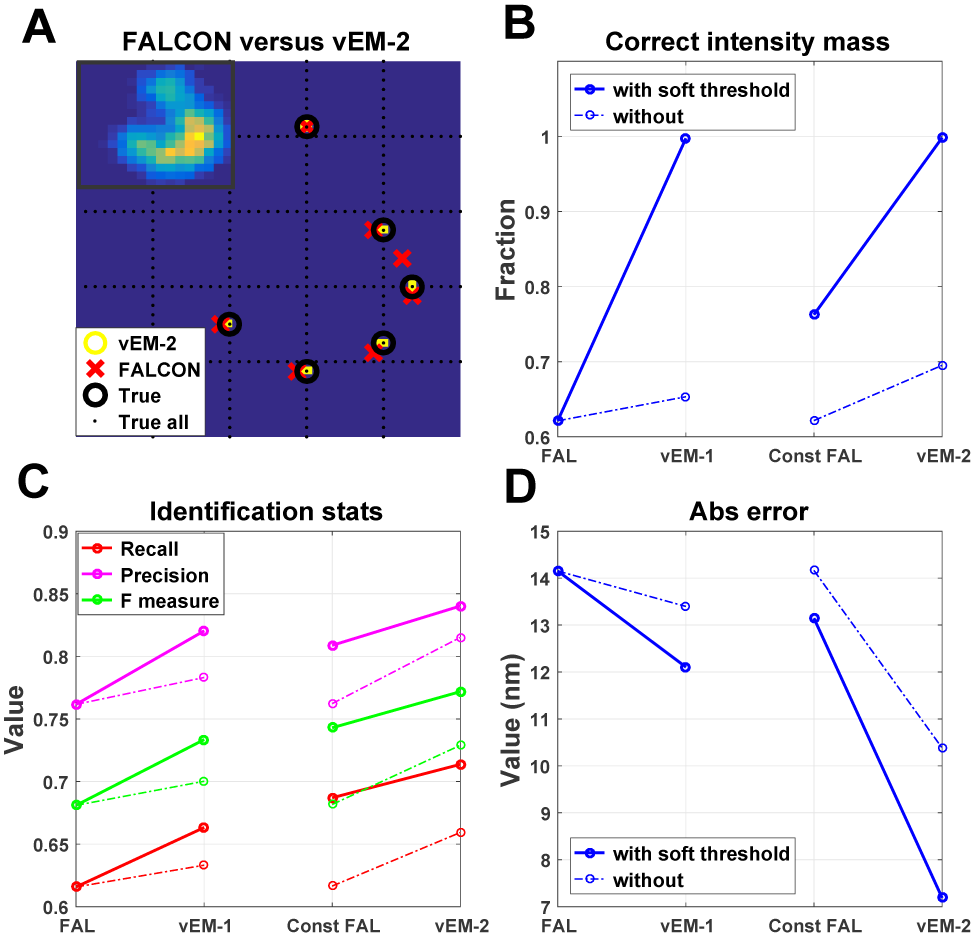
Evaluating performance of each algorithm step. (A) Estimates of active fluorophore locations in a single frame. Dotted black line indicates location of remaining (inactive) fluorophores. Note that vEM-2 estimates are more accurate than FALCON estimates. Algorithm steps and all simulation details follow Fig. 3, except in the vEM-1 step we compute results over all 5000 frames (not just 2000 frames, as in Fig. 3 upper right), for apples-to-apples comparison against the vEM-2 results here. Inset: observed data *Y*_*i*_ for this frame. **(B)** The percentage of 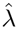 contained within the true support after each algorithm step. Note that vEM leads to significant improvements over FALCON; applying soft-thresholding in the M-step also provides significant improvements. **(C)** Recall, Precision and F-measure of identified fluorophores (solid: soft thresholded; dashed: no soft thresholding), and **(D)** mean absolute error of fluorophore location estimates. In both cases, similar trends as in **(B)** are visible.

Figure 5 provides a visual summary of one of the critical points of this paper: as the number of observed frames *N* increases, the vEM estimator continues to improve, and by *N* = 5000 is able to recover the true support of the underlying grid image with almost perfect accuracy. FALCON, on the other hand (as well as other approaches that estimate each frame independently), outputs estimates that appear blurry, due to noise in the estimated fluorophore locations, averaged over many frames – and this effective blur (and resulting loss of resolution) does not decrease asymptotically as N increases, since unlike vEM, FALCON does not exploit information from the (*N* – 1) other frames to improve estimation of individual frames.

**Figure 5:**
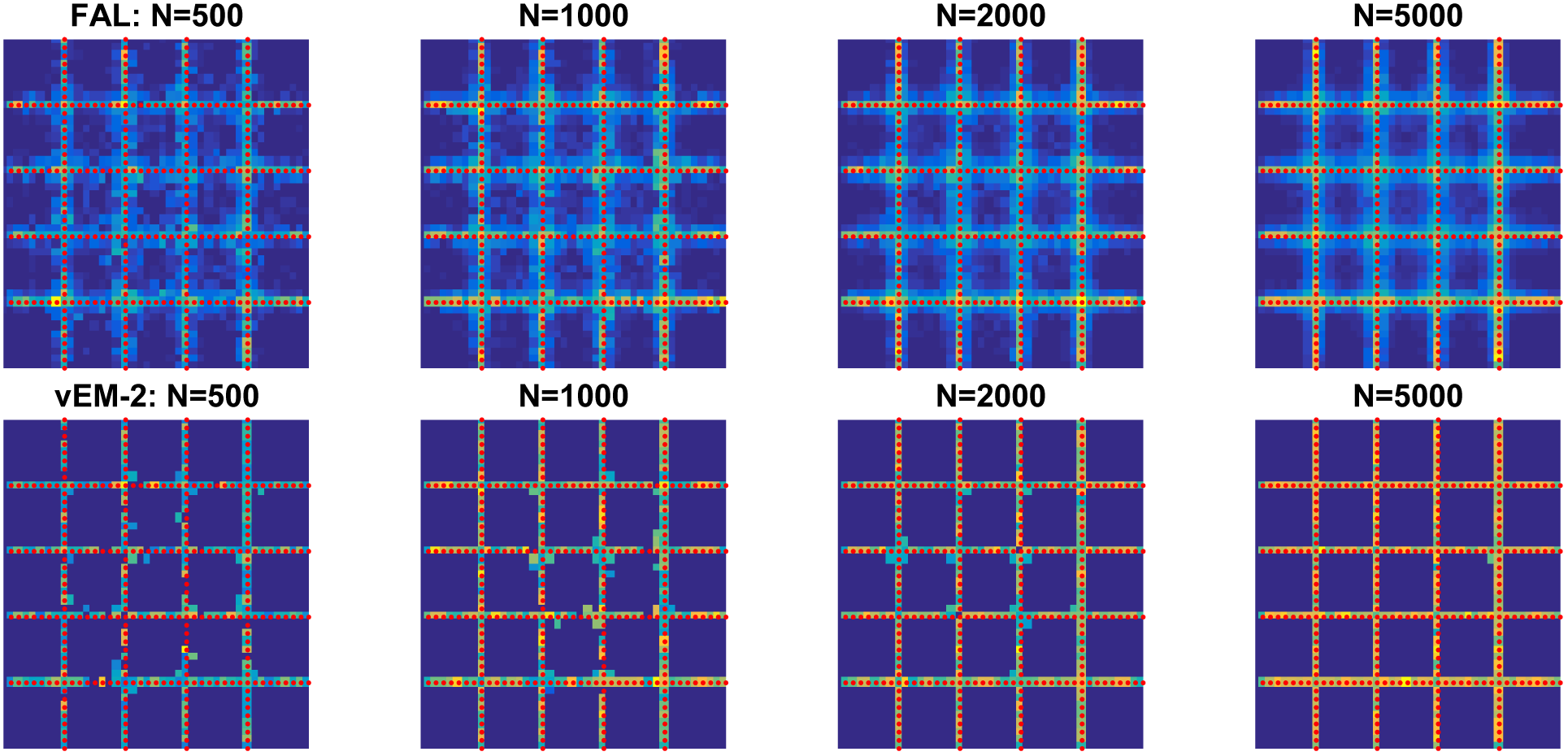
Estimates with varying number of observed frames N. Estimated 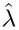 images output by FALCON (upper panels) and vEM (lower panels) with 500, 1000, 2000, and 5000 simulated frames (with imaging parameters such as PSF width and fluorophore density *p* held fixed over all frames). Red dots indicate the ground truth grid. The grid recovery accuracy of the vEM algorithm continues to improve with *N* – recovering the underlying grid structure nearly perfectly when *N* = 5000 – but the estimation noise-induced blur in the FALCON estimate does not decrease with *N*.

Figure 6 quantifies these results further, and adds comparisons to other competitive algorithms in the literature. Again the conclusion is that the vEM approach provides significantly more accurate estimates at little computational cost. Supplementary Figures 9 and 10 in the appendix show that this conclusion holds fairly uniformly over a wide range of PSF widths and average fluorophore densities, respectively.

**Figure 6:**
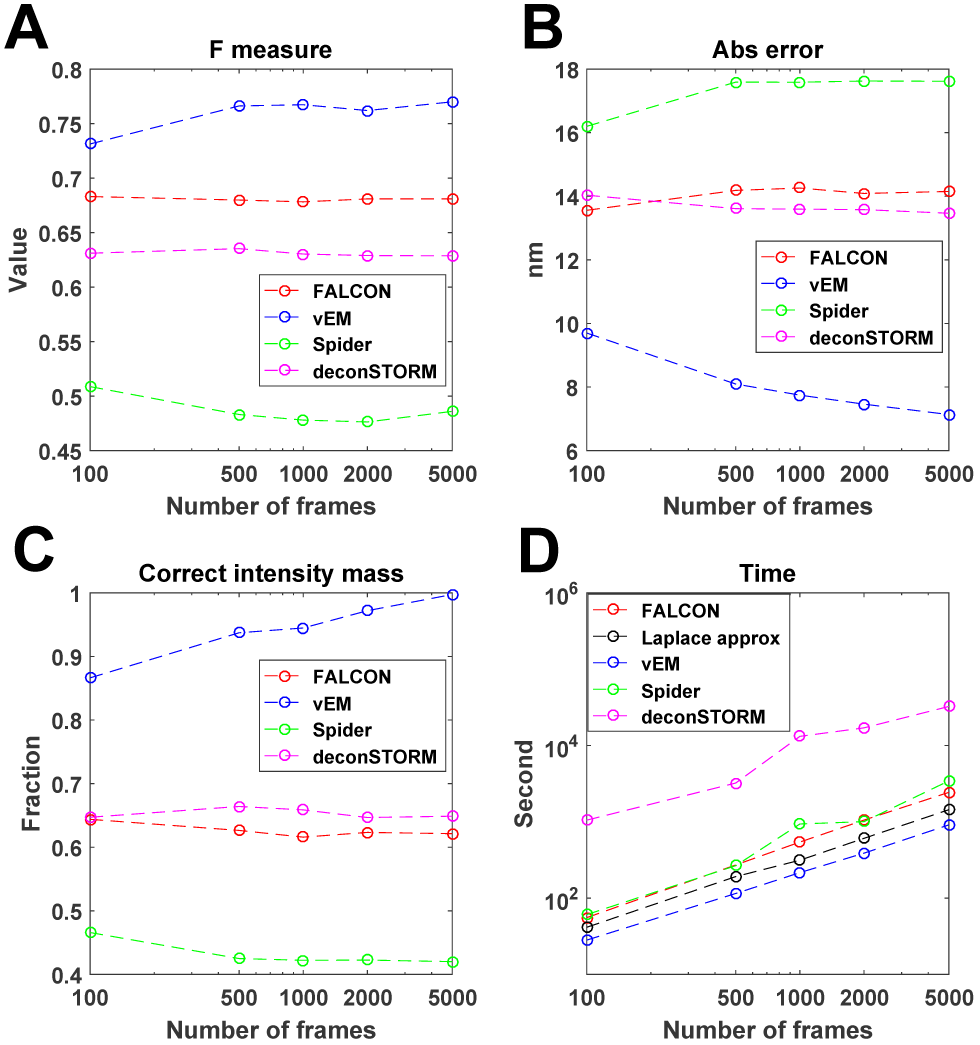
Evaluation of vEM in comparison with FALCON. [4], deconSTORM [6], and SPIDER [16] as a function of the number of observed frames N. A-C: F-measure, mean absolute fluorophore estimation error, and fraction of fluorophore mass recovered correctly on the true underlying grid computed as in Fig. 4. The vEM approach outperforms the other state-of-the-art algorithms on all of these metrics. D: Computational time of each algorithm step. Our full algorithm runs FALCON (red curve), computes the Laplace approximation (black curve), then iterates vEM to convergence (blue curve), then repeats the whole process on at least a subset of frames, so overall speed is ∼ 2x slower than FALCON overall. The deconSTORM algorithm is relatively much slower here.

Finally, Figure 7 shows a comparison of FALCON vs vEM applied to real data (see Appendix for full details). In this case the ground truth image is not available for comparison, but nonetheless the results are consistent with the simulated results described above: vEM leads to a sharper, better-resolved image than does FALCON.

**Figure 7:**
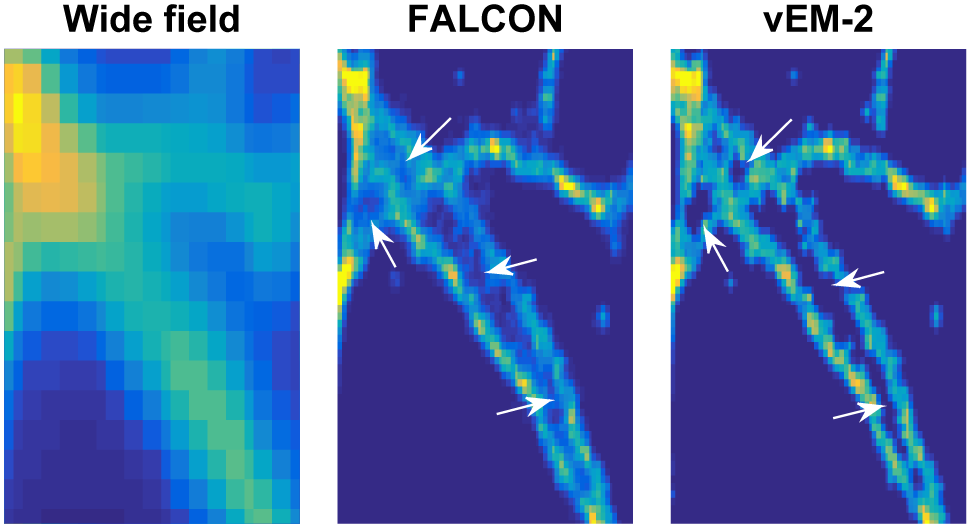
Analysis of real tubulin image data. Final resolved images output by FALCON and vEM-2 with 5000 frames. Wide field image, with all fluorophores turned on simultaneously, is given in first panel. Note that FALCON image is blurrier than the vEM-2 image, especially in areas of high fluorophore density, e.g., where multiple tubulin branches are close together, as noted by white arrows.

**Figure 8:**
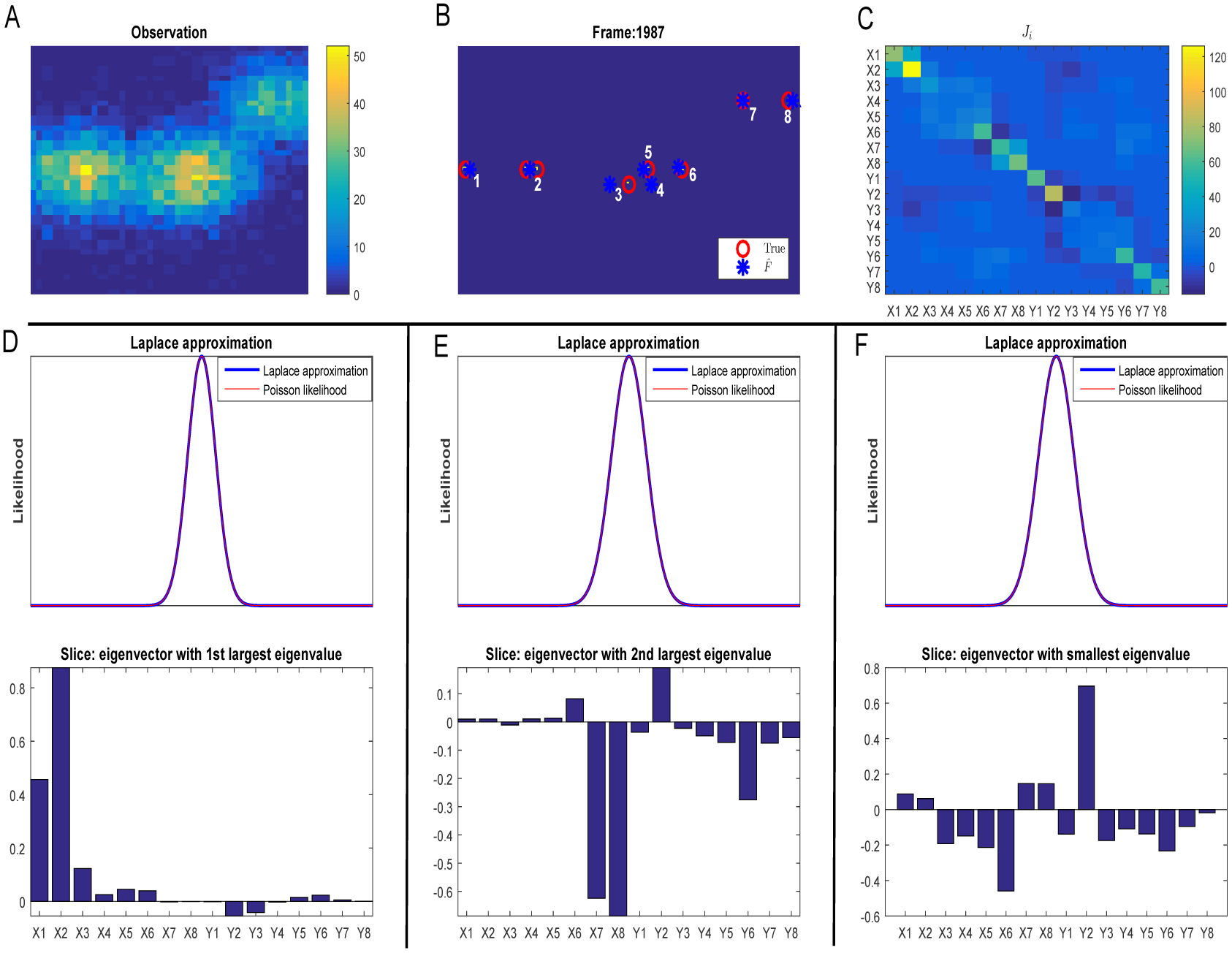
Accuracy of the Laplace approximation. We use the same frame as in Figure 2 in the main text as an illustration. (**A**) *Y*_*i*_. (**B**) Red circles are true positions; blue stars are 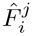. (**C**) Fisher information matrix *J*_*i*_. Xj indicates *x* coordinates of fluorophore *j*, and Yj the *y* coordinates. FALCON inferred 8 fluorophores in this frame, so we have 16 total coordinates. Note that the blocks of *J*_*i*_ corresponding to fluorophores 3,4,5,6 have smaller values, indicating reduced estimation accuracy due to overlapping PSF bumps. Panels (**D**), (**E**), (**F**) indicate the accuracy of the Laplace approximation for the Poisson likelihood. Since the likelihood w.r.t. *F*_*i*_ is a 16-dimensional function, we can only display slices of this function. We choose slices in the direction of eigenvectors of *J*_*i*_ – the principal components of the Laplace approximation. (Directions appear in bottom panels.) The red (Poisson likelihood) and blue (Laplace approximation) curves align very closely in each of the three directional slices shown here.

**Figure 9:**
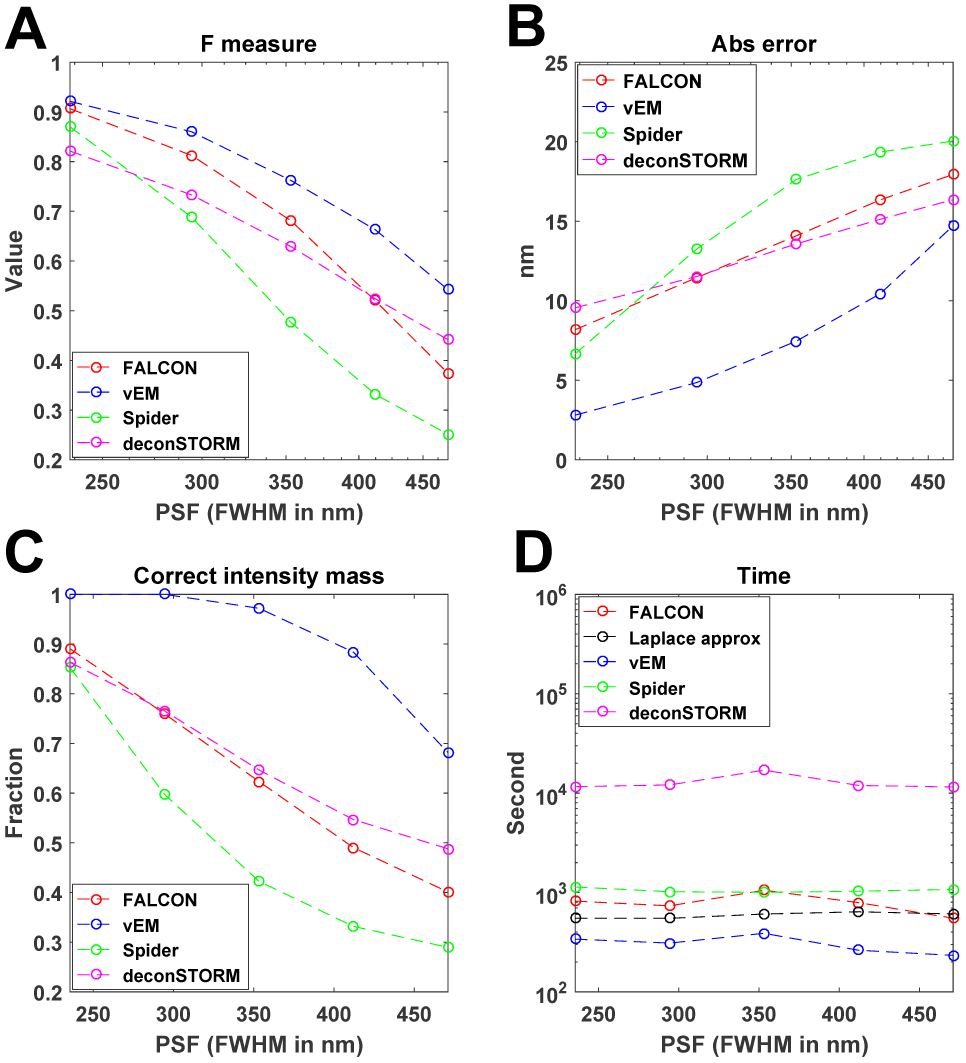
Evaluation of vEM in comparison with FALCON. [4], deconSTORM [6], and SPIDER [16] as a function of PSF width. Panel layout as in Fig. 6. 2000 frames were used here.

**Figure 10:**
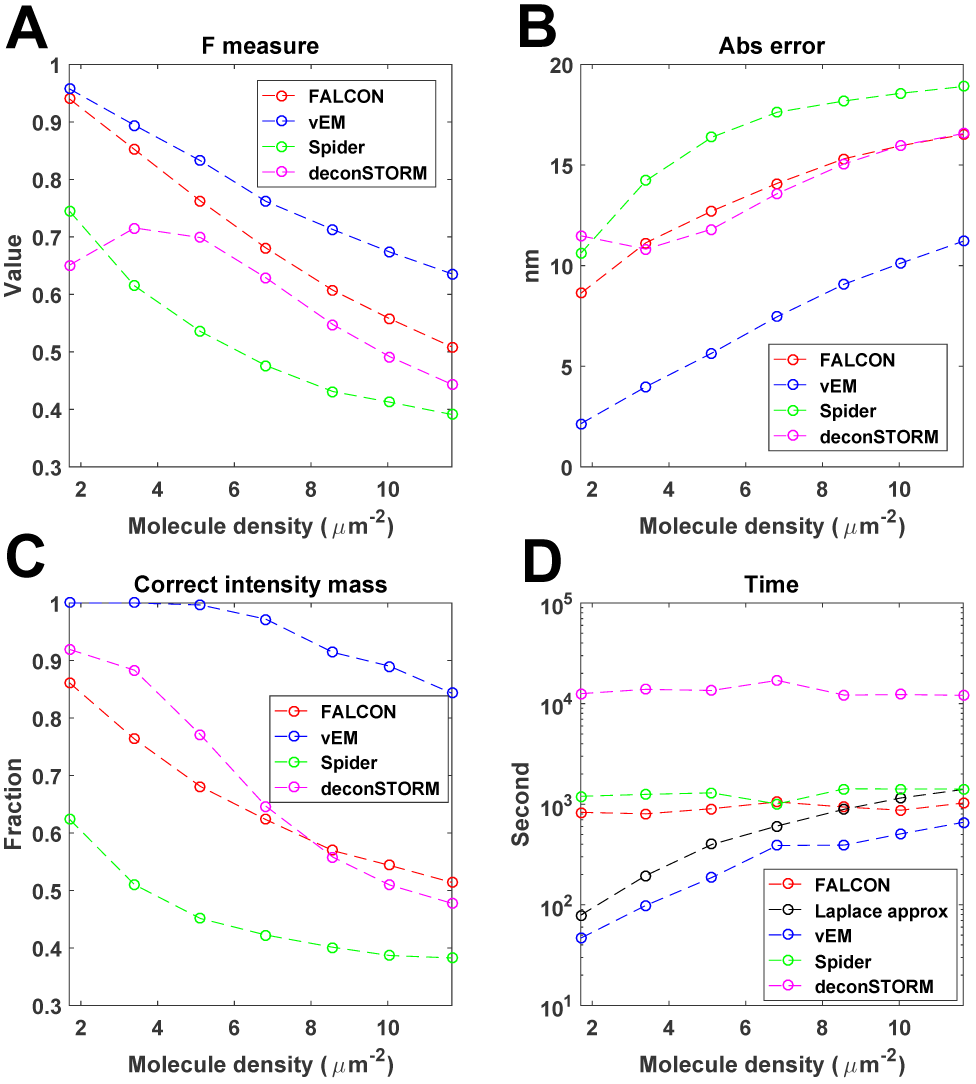
Evaluation of vEM in comparison with FALCON [4], deconSTORM [6], and SPIDER [16] as a function of fluorophore density. *p*. Panel layout as in Figures 6 and 9. We found that deconSTORM tends to have an inflated false positive rate (i.e., lower precision) for small values of p in panel (A). Also note that vEM is relatively cheaper computationally for small *p*, where there are fewer fluorophores to iterate over in the Laplace approximation and vEM steps. 2000 frames were used here.

## Discussion

We have introduced scalable Bayesian methods for improved estimation in super-resolution microscopy. By further extending the reach of these critical imaging methods, our approach can significantly impact a variety of biological applications. The hybrid vEM / Laplace-approximation / sparse-representation approach developed here is more generally applicable in other hierarchical sparse signal model applications [11]. Our methods exploit the insight that sharing information across image frames significantly improves accuracy – and this effect grows more powerful as the number of frames *N* increases.

Similar points have appeared previously in the super-resolution microscopy literature, notably in [6] and [7]. The methods introduced in [7] are seldom used in practice on large-scale imaging data, due to prohibitive computational expense. The vEM methods we have introduced here are much more scalable (Fig. 6D); indeed, we were unable to obtain good results from the method used in [7] in a reasonable amount of computational time (> 1 day) and so we did not show comparisons against this method here (see appendix for further discussion).

The deconSTORM method described in [6] (see also Fig. 6) attempts to improve upon simple Richardson-Lucy deconvolution by incorporating local information about the survival of active fluorophores from one frame *i* to the next (*i* + 1). Our approach is orthogonal: we share information between frames *I*_*i*_ *globally*, through 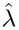. As we discuss in the appendix (“Markov model”), the vEM framework extends easily to handle these local correlations between fluorophores at frames *i* and *i* + 1. We observed that although incorporating these local correlations can slightly improve the recovery of individual fluorophores (Fig.11), the local Markov model does not significantly qualitatively improve the accuracy of the final estimated 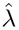 (Fig.12).

**Figure 11:**
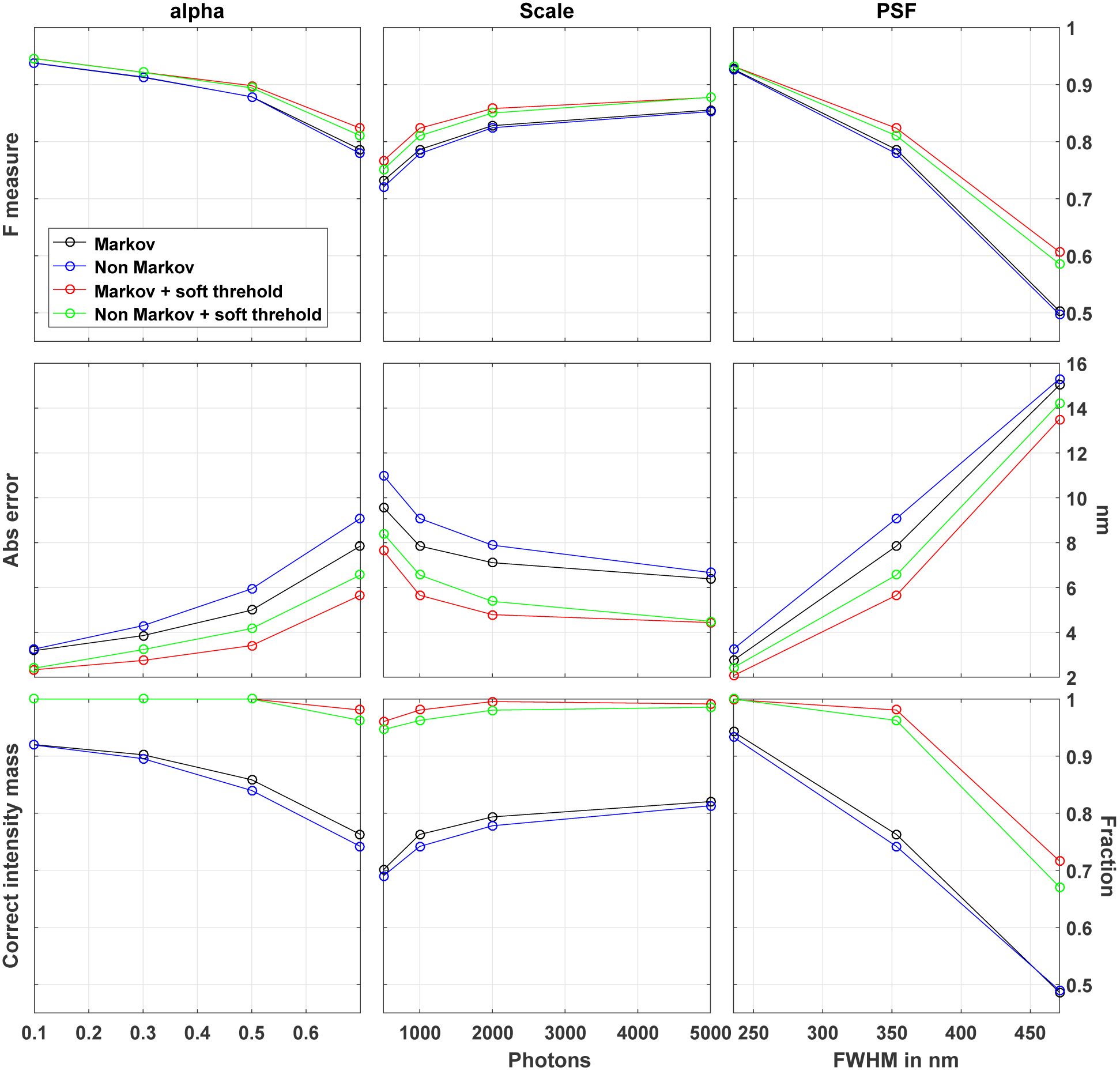
Evaluation of Markov and non-Markov models as a function of *α* (First column), *Scale* (Second column), and *PSF* (Third column), with and without soft threholding. The performance is quantified using the same measures as in the main text. *Scale* is the average number of photons per active fluorophore. We use *N* = 2000 frames. Emission rate *p* is 0.01. When not indicated otherwise, *α, Scale,* and *PSF* are set to be 0.7, 1000 photons, and 353.25 nm respectively. Note that this is rather large value of *α* as seen in the left panel, smaller differences between the Markov and non-Markov models are seen when smaller values of *α* are used.

**Figure 12:**
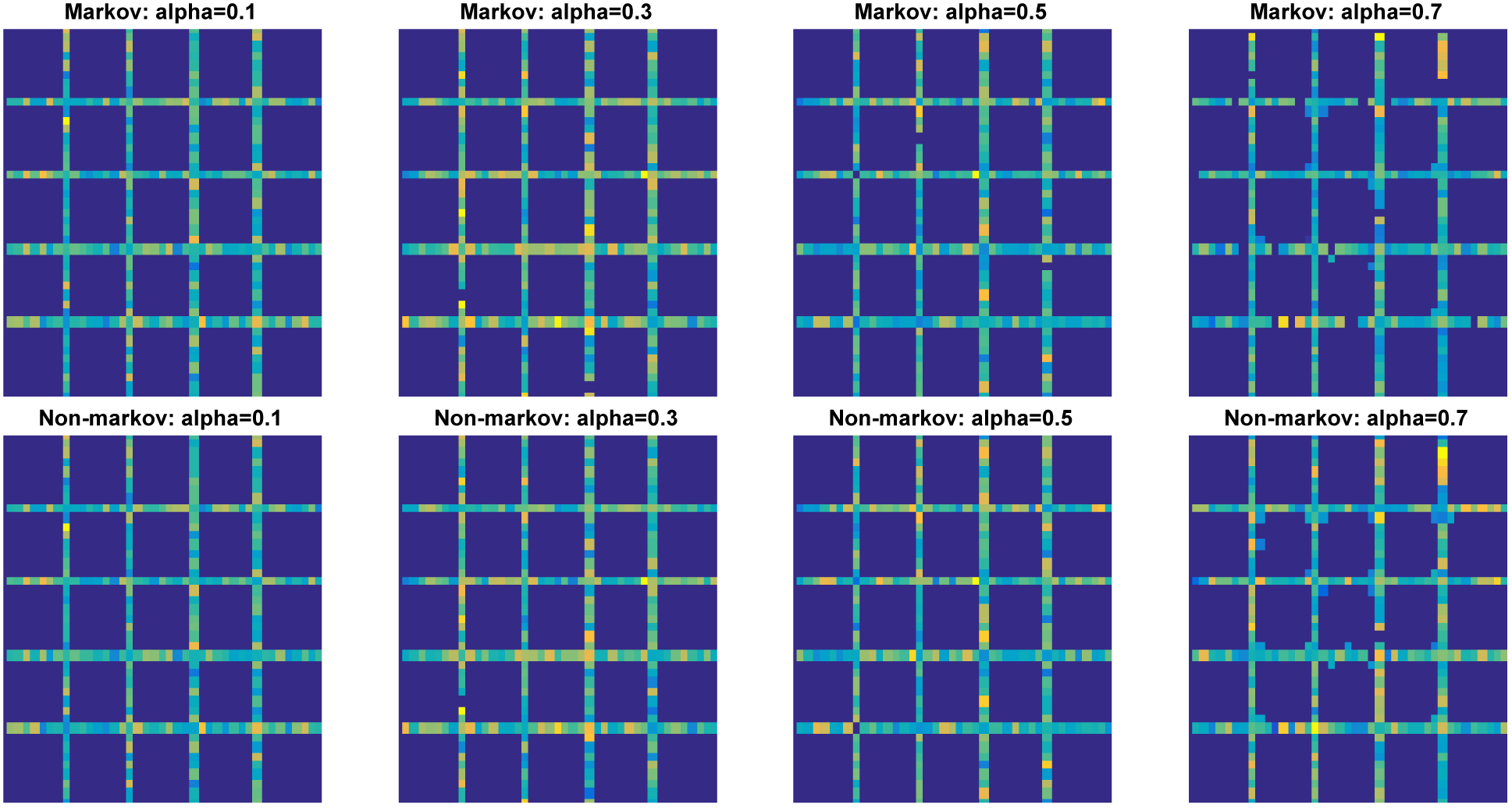
Estimates of Markov and non-Markov model as a function of *α* with soft thresholding̤. The final resolved images of Fig. 11 (First column). The test image was the same grid used in the main text. Recall that *α* influences the number of active fluorophores (with large *α* corresponding to more persistent fluorophores and therefore higher fluorophore density); the average fluorophore densities in the four columns shown here are 1.90, 2.55, 3.44, and 5.75 *µ*m^−2^ (left to right). We use *N* = 2000 frames. Emission rate *p* is 0.01, with an average of 1000 photons per fluorophore. PSF is 353.25 nm. Note that no major differences are seen between the Markov and non-Markov estimates. Similar results were obtained over a wide range of parameters (not shown).

A final interesting and important direction for future work would be to extend some of the methods developed here to the case where the fluorophores are moving from frame to frame, in the context of single-particle tracking experiments [17].

## Acknowledgments

We thank Professor John P. Cunningham for helpful discussions, and Ido Rosen for helpful comments on our code. Funding for this research was provided by Simons Foundation Global Brain Research Award 325398, ONR N00014-14–1-0243, ARO MURI W911NF-12–1-0594, and a Google Faculty Research award; in addition, this work was supported by the Intelligence Advanced Research Projects Activity (IARPA) via Department of Interior/ Interior Business Center (DoI/IBC) contract number D16PC00008. The U.S. Government is authorized to reproduce and distribute reprints for Governmental purposes notwithstanding any copyright annotation thereon. Disclaimer: The views and conclusions contained herein are those of the authors and should not be interpreted as necessarily representing the official policies or endorsements, either expressed or implied, of IARPA, DoI/IBC, or the U.S. Government.

## APPENDIX “Scalable variational inference for super resolution microscopy”

## Details of algorithmic comparisons

All parameters of the methods compared here are tuned for best performance.

### FALCON [4]

We set the sparsity parameter (the *ℓ*_1_ weights) *κ* to 3. The threshold above which the support is defined is set as 10% of the maximum intensity.

### SPIDER [16]

We set the sparsity parameter, the weights of *ℓ*_*o*_, regulation, *κ* = 250.

### deconSTORM [6]

In our simulations accuracy continued to improve even after 5000 iterations, so we used 5000 iterations in our comparisons. Note computation time is proportional to the number of iterations, so this method could be sped up at the cost of some accuracy.

### 3B [7]

As mentioned in the Discussion, we were not able to obtain reasonable results using this method, even after > 1 day of computation, for data sets with e.g. 2000 observed frames. In personal communications with the developers of this method, it was emphasized that this approach is better suited for smaller images and smaller values of *N*, since the speed of this method decreases more or less with the square of the size of the PSF (in pixels), the size of the image area being analyzed, and the number of frames observed. Therefore we did not pursue further quantitative comparisons against this method.

### Evaluation

Identified fluorophores are defined as estimates within some fixed distance from the true fluorophores. The cutoff radius we used was 50 nm. FALCON returns a list of fluorophore locations, whereas vEM, deconSTORM, and SPIDER return images of the estimated fluorophore density. To quantify fluorophore estimate accuracy for these methods we thresholded these images and took local weighted averages to obtain the estimated fluorophore locations.

## Experimental details

### Simulation

To validate our analysis method, we simulated grid data on a 32-by-32 pixel map with an image pixel size of 100 nm. The final resolved image sits on a 3x finer grid with super resolution pixel size of 33 nm. Unless stated otherwise, each frame has an emission rate of 0.04, corresponding to an average molecule density of 6.8 *µm*^−2^. The average photon number is 1,000 per fluorophore with PSF width 150 nm in standard deviation or 353 nm in FWHM. To test the performance of our method under different practical situations, we varied critical parameters, including the number of frames, molecule density, and PSF width. In those simulations, we replace the above parameters with a range stated in the corresponding figure caption and keep other parameters unchanged.

### Real data

We reconstruct a patch of tubulins on a 32-by-32 pixel image map with 5000 real experimental frames. The final resolved image sits on a 4x finer grid. We modeled the PSF as a Gaussian blur with width of 183 nm in standard deviation; this parameter was estimated by fitting observations of nonoverlapping fluorophores to a 2D Gaussian function, following [18]. The dataset is provided as Tubulin ConjAL647 on the Single Molecule Localization Microscope website [3].

### Markov model

In the main text, we focus on a model in which fluorophores become active according to a Poisson process with rate λ. The active fluorophores in one frame are conditionally independent from those in other frames, given λ. In this section, we incorporate the phenomenon that some fluorophores do not quench immediately, i.e. there is a probability that an active fluorophore will remain active in the following frames. Thus the active fluorophores in one frame consist of two groups: “newborn” fluorophores that activate from the dark state with rate λ (the same as before), plus fluorophores remaining active from the previous frame, each with probability *α*. Under this assumption, the active fluorophores in one frame are dependent on those in the frame before and after, so we denote the new model as the “Markov model” and the original model as the “non-Markov model.”

The Markov model incorporates the positions of active fluorophores in neighboring frames *i* + 1 and *i* – 1, which can potentially be useful to help pinpoint the positions of active fluorophores in each frame *i.* Thus we would expect the Markov model to outperform the non-Markov in localizing individual fluorophores. (Similar neighboring-frame effects are incorporated in [6] and [7].) In this section, we will first introduce the Markov model, and then show results comparing the effectiveness of the Markov model versus the non-Markov model.

### 1 The model

We begin by writing down the Markov model for the time series of activations of fluorophores *I*_:,*xy*_ at location *xy* across all *N* frames:

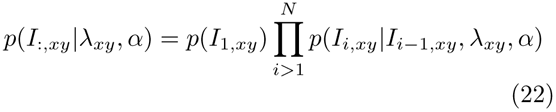

(As usual, fluorophores in different locations *xy* activate conditionally independently given λ.) The transition matrix is given by

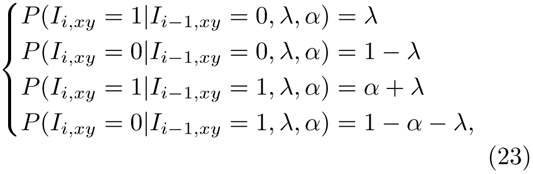

where *α* is the probability that an active fluorophore remains active in the next frame, and λ is defined in eq. 2, 3; note that the probability of a new fluorophore activating in any frame is typically fairly low (to guarantee that each image *I*_*i*_ is sparse), so 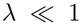, while the probability of remaining active may be non-negligible. We are most interested here in the case that *α* is significantly greater than 0, implying 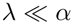. Finally, note that we use a Bernoulli emission model here instead of the Poisson emission model used in the main text (eq. 5); this simplifies the derivations below. Of course the Poisson and Bernoulli models are identical in the limit of small λ.

After the Laplace approximation our full approximate loglikelihood is

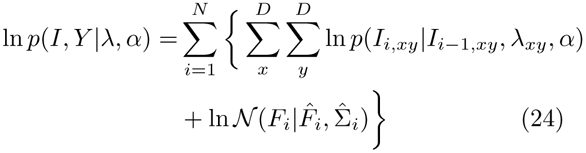

where in eq. 24 we define *I*_0_ as all zeros.

### 2 Variational EM Algorithm

Now we proceed as before and maximize the ELBO (eq. 9) to obtain the E and M steps. We will assume that *α* is known. In reality, of course, *α* is unknown and needs to be estimated along with λ. It is straightforward to derive EM iterations for *α* as well, but we do not pursue this here. Instead, in the Results section we will simulate data from the Markov model and estimate λ using a known value of α, to give the Markov approach the best possible chance of improving over the results of the non-Markov model.

#### M step

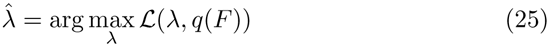

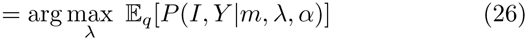

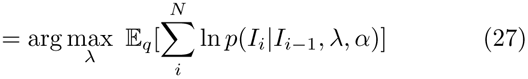

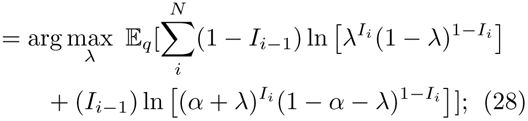

eq. 28 combines the cases in eq. 23 and uses the property that *I*_*i*_ can only take values of 0 or 1. We have dropped the pixel subscript *xy* to simplify notation; the maximization problem is separable over locations *xy* and therefore we can optimize for each pixel independently in parallel.

Setting the derivative w.r.t. λ to zero, we obtain

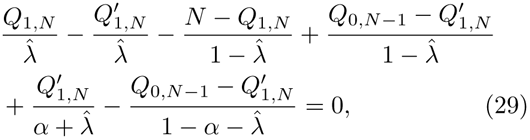

where we define 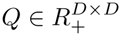 and 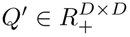

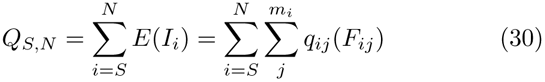

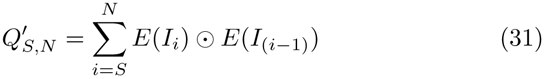

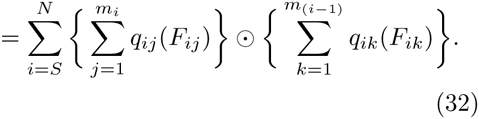

Now set

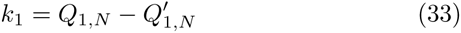

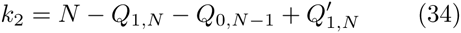

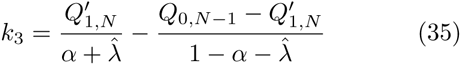

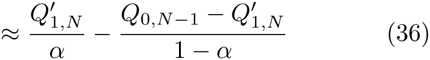

where in eq. 36, we used λ ≪ α. Therefore, eq. 29 becomes

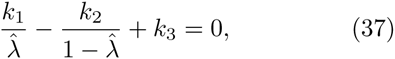

which reduces to a simple quadratic equation in 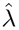. We select the valid solution 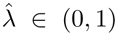. Similarly as in the non-Markov model, we can perform soft thresholding on the obtained value to increase the sparsity of 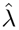.

Finally, note that if we set *α* = 0, eq. 29 becomes

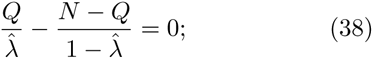

this leads to 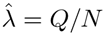, which corresponds to the M step in the non-Markov model (description under eq. 16), as desired.

#### E step

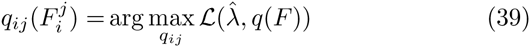

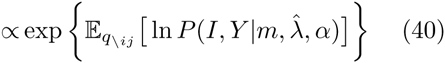

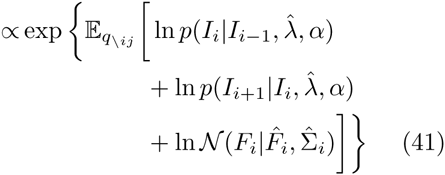

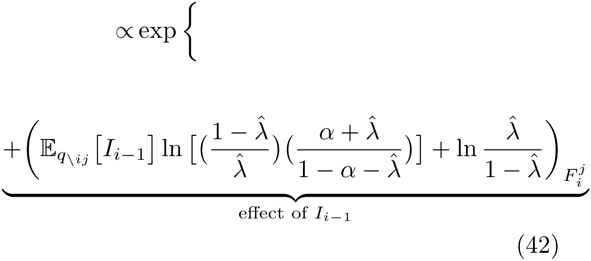

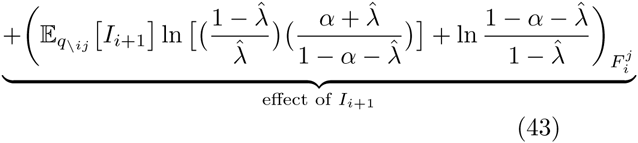

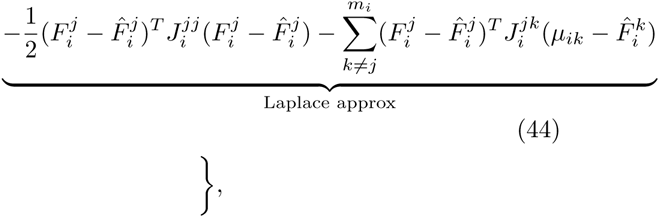

where the operator 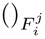 is the value of the 2D function of the variable in the parentheses at location 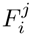.

Note that if *α* = 0 and 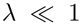, then

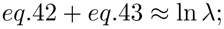

therefore, we recover the E step in the non-Markov model (eq.20), as desired.

### 3 Results

To quantify the benefits of including the Markov terms in the model, we generate data from the Markov model with a range of parameters and perform inference with both the Markov and non-Markov models. Note that the non-Markov model is mis-specified in these simulations, while the Markov model is given the “unfair” advantage of knowing the true value of *α*. Nonetheless, somewhat surprisingly, our basic conclusion is that incorporating the Markov effects has only a small effect on inference performance. Fig. 11 shows that the estimation accuracy on individual fluorophores is improved modestly if the Markov terms are included, once *α* is sufficiently large. However, in Fig. 12 we see that the Markov terms lead to negligible improvement in the overall estimate of λ, which is the main object of interest in many super-resolution imaging studies. Thus we conclude that the “local” information encoded by the Markov terms in the model is mostly redundant with the “global” information encoded by our estimate 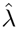, which is shared across frames to improve our estimate of each *I*_*i*_.

